# Stable and functionally diverse versatile peroxidases by computational design directly from sequence

**DOI:** 10.1101/2021.11.25.469886

**Authors:** Shiran Barber-Zucker, Vladimir Mindel, Eva Garcia-Ruiz, Jonathan J. Weinstein, Miguel Alcalde, Sarel J. Fleishman

## Abstract

White-rot fungi secrete a repertoire of high-redox potential oxidoreductases to efficiently decompose lignin. Of these enzymes, versatile peroxidases (VPs) are the most promiscuous biocatalysts. VPs are attractive enzymes for research and industrial use, but their recombinant production is extremely challenging. To date, only a single VP has been structurally characterized and optimized for recombinant functional expression, stability and activity. Computational enzyme optimization methods can be applied to many enzymes in parallel, but they require accurate structures. Here, we demonstrate that model structures computed by deep-learning based *ab initio* structure prediction methods are reliable starting points for one-shot PROSS stability-design calculations. Four designed VPs encoding as many as 43 mutations relative to the wild type enzymes are functionally expressed in yeast whereas their wild type parents are not. Three of these designs exhibit substantial and useful diversity in reactivity profile and tolerance to environmental conditions. The reliability of the new generation of structure predictors and design methods increases the scale and scope of computational enzyme optimization, enabling efficient discovery and exploitation of the functional diversity in natural enzyme families.

## Introduction

The need to develop economical and environmentally friendly energy sources is undeniable. Efficient conversion of biomass, particularly lignocellulose, into biofuels is a promising route for sustainable and renewable energy production^1^. The amorphous and highly cross-linked structure of lignin, however, obstructs the accessibility of chemicals and enzymes to cellulose and impedes their conversion into biofuels and other high-value chemicals^2–4^. Furthermore, lignin itself comprises potentially valuable chemicals that could be valorised. Since chemical depolymerization of lignin is still not economically viable or environmentally benign^2,5^, biodegradation is an attractive route for utilization of wood biomass.

The most efficient natural system for lignin depolymerization is observed in white-rot basidiomycetes. These fungi secrete a repertoire of high-redox potential oxidoreductases (laccases and peroxidases) which degrade lignin synergistically^6–8^. Of those, versatile peroxidases (VPs; EC 1.11.1.16) are of particular interest for biotechnological use due to their broad substrate scope ranging from low to high-redox potential substrates. VPs reduce hydrogen peroxide by oxidizing a wide range of substances, including phenolic and non-phenolic compounds, pesticides, high-redox potential dyes, polycyclic aromatic hydrocarbons and lignin^9^. Some fungal species secrete several VP paralogs, suggesting that VPs may act synergistically^10,11^. Nevertheless, VPs are especially challenging for production in heterologous hosts, limiting their use in research, let alone as an enzyme repertoire or in industrial applications.

One reason why VPs are functionally promiscuous is that they comprise three distinct active sites for substrate oxidation: a site for the oxidation of Mn^2+^ to Mn^3+^, which acts as a diffusible mediator, a low-redox potential heme-dependent binding pocket, and a high-redox potential surface reactive tryptophan radical which connects to the heme through a long-range electron transfer pathway^9^. Additionally, they comprise two structural calcium ions, multiple glycosylations, and several disulfide bonds, thereby complicating their expression in heterologous hosts. Thus, to date only a VP from *Pleurotus eryngii* (VPL) has been fully characterized at the biochemical and structural levels^12^. Several directed evolution campaigns successfully adjusted VPL to various industrial requirements: functional expression in the yeast *Saccharomyces cerevisiae*, thermostability^13^, stability and activity in neutral and alkaline pH^14^, and stability and activity under high concentrations of H_2_O_2_^15^ which serves as the terminal electron acceptor in VPs but is also a strong inhibitor. In each such campaign, 5,000-15,000 clones generated by random mutagenesis and diverse DNA recombination methods were screened to reach the desirable trait^13–15^. Although successful, the high labor intensity makes directed evolution an impractical approach for optimizing multiple natural starting points. In such cases, computational design methodologies may provide a useful alternative^16^. The PROSS structure-based algorithm combines phylogenetic sequence information with Rosetta atomistic modeling to incorporate stabilizing mutations and optimise the native-state energy^17,18^. As such, PROSS has addressed complicated protein-expression and stability problems by experimental screening of only a few designs (typically ≤ 5). In a number of cases, proteins that could not be functionally expressed in microbial systems could be expressed robustly after one-shot design calculations^17–19^. PROSS is sensitive to atomic details, however, and except in a handful of cases^20,21^, has not been applied to structural models.

Recently, a new generation of deep-learning-based *ab initio* structure prediction methods has been developed, with the most recent ones reaching the accuracy of crystal structures^22,23^. Here, we introduce a general pipeline for increasing the functional expression yields of proteins for which no experimental structure data are available using these structure predictors. The structural and biochemical complexities of VPs make them especially challenging as a proof-of-concept for this pipeline. First, we selected 11 diverse VP sequences, modeled them using the trRosetta structure prediction algorithm made available in December 2020^24,25^ and ran PROSS^17,18^ stability-design calculations on each as well as on VPL (in total 12 natural VPs; Figure 1). After screening only 36 designs (three for each wild type VP), we isolated three efficient, highly stable VPs that can be functionally expressed in yeast whereas their wild type progenitors could not be recombinantly expressed efficiently. Our approach exploits nature’s diversity to generate a VP repertoire that can be used for decomposition of lignin and diverse pollutants. Furthermore, the combination of the structural and experimental data we obtained expands our understanding of VP structure-function relationship.

**Figure 1.**
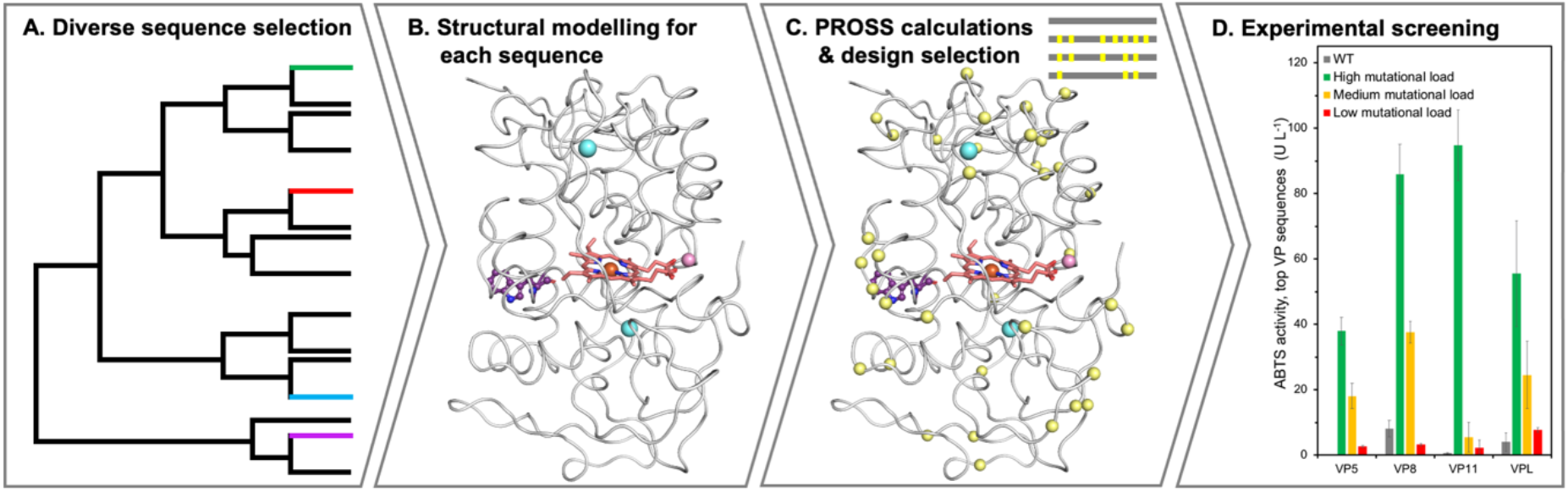
Key steps in the design of a diverse set of VPs. (A) VP sequences were collected from different databases, a phylogenetic tree was computed and twelve sequences that represent all branches were selected for diversification. (B) The selected sequences were modeled by trRosetta^24,25^. For visualization, heme (red), manganese (pink) and calcium ions (blue) were superimposed from the VPL structure (PDB entry: 3FJW). The surface-reactive tryptophan is presented in purple balls-and-sticks. (C) PROSS stability design calculations^17,18^ suggested dozens of mutations (yellow spheres). For each sequence, three designs with different mutational loads were selected for further experimental examination. (D) In an activity screen of proteins heterologously produced in yeast, the wild type proteins show negligible functional expression while the designs with the highest mutational load are highly active on the peroxidase substrate ABTS.

### Computational design for functional expression directly from sequence

VP sequences were extracted from public databases, phylogenetically classified, and eleven sequences that exhibited 51-81% identity to one another were selected for modeling using trRosetta (Figure 1A & B; Table S1 & S2; see SI *Methods* for more details)^24,25^. The top model in each case was then subjected to PROSS stability design (Figure 1C)^17,18^. VPL was also designed by PROSS based on its crystallographic structure (PDB entry: 3FJW). The new structure prediction methods do not model ligands and ions. We visually compared the models to the VPL experimental structure finding that the models retained the intricate arrangement of amino acids in the heme binding pocket and the ion-binding sites (Figure S1). This observation encouraged us that the trRosetta models could be used as reliable starting points for the design of enzymes as complex as the VPs. Based on the VPL crystallographic structure^26^, we inferred which positions in each model comprised the catalytic and ion-ligand sites and disabled design in these positions. We also restricted design in positions where model uncertainty was high and in adjacent positions (see SI *Methods*). Finally, we selected the wild type sequence and three PROSS designs with different mutational loads for experimental characterization (approximately 10 mutations in the most conservative design and up to 43 in the most permissive one, see Table S1; VPs comprise approximately 340 amino acids, *i.e.* roughly 12% of the protein template was mutated in the most permissive designs). The DNA encoding each protein was codon optimized for yeast expression, ordered as synthetic gene fragments that were incorporated in the pJRoC30 plasmid downstream of the *S. cerevisiae* α factor prepro-leader, and transformed into yeast cells.

The approach described above yielded four diverse and functionally expressed VPs (Figure 1D): VPL (from *Pleurotus eryngii*), two paralogs from *Pleurotus ostreatus* (VP5 and VP11) and a VP from *Ganoderma* sp. 10597_SS1 (VP8) (Table S3). For these VPs, the wild type progenitors demonstrated poor functional expression while the designs efficiently oxidized the peroxidase substrate 2,2’-azino-bis (3-ethylbenzothiazoline-6-sulfonic acid) (ABTS) (Figure 1D). Particularly, for all four VPs the most active design exhibits the highest mutational load (38-43 mutations, > 10% of the sequence); the designs are therefore denoted as 5H, 8H, 11H, and VPLH to designate their high mutational load. The designed mutations exhibit improved core packing, introduce new hydrogen-bond networks, and rigidify loops (Figure S2). Some of these mutations are radical (for example, an Ile→Phe mutation in the protein core; Figure S2A) and in many cases, proximal mutations form multiple new contacts, demanding atomic accuracy in the starting models.

### Designed VPs are highly stable

Since the wild type VPs showed negligible functional expression, all further biochemical analyses were done on 5H, 8H and 11H relative to the VPL variants R4 and 2-1B (previously evolved for expression and thermostability, respectively)^13^. Although VPLH also demonstrated significantly enhanced functional expression compared to its wild type protein (Figure 1D), it did not show substantial improvement relative to R4 and 2-1B (which are also derived from VPL) and was not pursued further.

The designed VPs exhibit higher thermal stability compared to R4 (Figure 2A & B and Figure S3). While the temperature at which the enzyme loses half of its maximal activity levels after 15 min incubation (T50) is similar to R4 across all designs, at elevated temperatures, 5H does not lose its activity completely (Figure S3A). In shorter incubations, 5H shows much enhanced activity at 45-60 °C compared to the activity at room temperature, similar to 2-1B, and the highest residual activity at 65-80 °C (Figure S3B). This trend is consistent with the observed kinetic stability at 60-65 °C (t1/2, the time at which the protein loses half of its activity after incubation at a specific temperature; Figure 2B and Figure S3C): 5H maintains stable residual activity even after two hours, comparable to 2-1B, which was evolved specifically to withstand high temperatures. Long-term heat resistance, as observed for these designs, is an important advantage when using enzymes in industrial processes.

**Figure 2.**
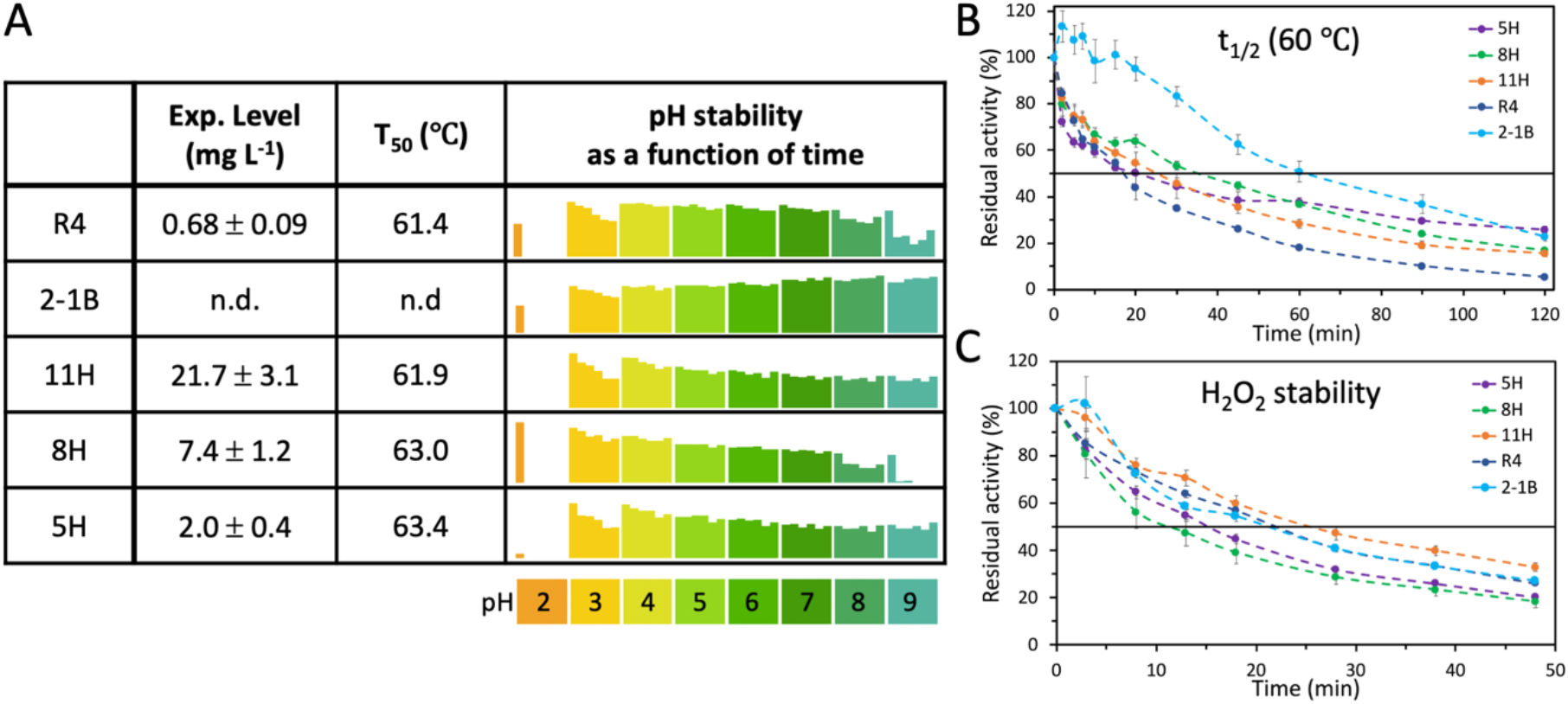
Functional expression levels, thermal, pH and H_2_O_2_ stability in VP designs. (A) The expression levels were calculated using the initial activity against ABTS in the supernatant immediately after growth and the ABTS kinetic values (see SI *Methods*). Apparent T_50_ values were calculated based on the residual activity of enzymes incubated at different temperatures (see Figure S3A); n.d. : not determined. pH stability was assessed by incubation at pH ranging from 2-9 and measuring the residual activity at times 0, 4, 24, 50, 75 and 165 hours, compared to the activity at pH=3 at time zero (see Figure S4 for complete data). (B) Kinetic thermostability (t_1/2_) profiles were determined by incubating VP supernatants at 60 °C and measuring their residual activity at times 0-120 minutes, compared to the initial activity. (C) H_2_O_2_ stability (t_1/2_) profiles were determined by incubating designs in H_2_O_2_ in a molar ratio of 1:3,000 and measuring their residual activity at times 0-48 minutes, compared to the initial activity. All the results are the means ± S.D. of three independent experiments.

In nature, lignin is decomposed under acidic conditions^11^, and VP activity is strongly acid-dependent^14,27,28^. Therefore, high oxidizing power depends on stability and activity at pH 2-3^14^. 8H is stable under acidic to neutral pHs and is the most active VP variant after incubation in highly acidic pH (pH=2; Figure 2A and Figure S4). VPs may also be useful in alkaline conditions, for instance for biomedical purposes and for paper and textile processing^8^. Remarkably, designs 5H and 11H maintain their initial activity levels even after one-week incubation at pH 9 (Figure 2A and Figure S4A).

Hydrogen peroxide is the terminal electron acceptor in VPs but at high concentrations it also deactivates enzymes^29^. To assess the stability at high hydrogen peroxide concentrations, VPs were incubated with H2O2 at 1:3,000 molar ratio (Figure 2C). Of all VP variants, 11H exhibits the highest stability to hydrogen peroxide.

Lastly, we estimated the expression levels of all VPs based on initial rate measurements in the yeast broth derived from screening experiments, and the kinetic constant data (see Table S4, SI *Methods* and below). Under these specific conditions which were not optimized for high protein expression, the three VP designs show much greater functional-expression levels than R4, which is the highest functionally expressed VPL variant^13^ (Figure 2A). Further gains in functional expression can be made by optimizing the experimental conditions or expression strain, as was previously noted in the R4 directed evolution campaign^13^. We concluded that the designs were stable and exhibited remarkable differences in their tolerance of environmental conditions.

### Designs exhibit diverse functional profiles

We next tested the activity profiles of the top-three VP designs with a range of peroxidase substrates using R4 as a reference (Figure 3, Figure S5 and Table S4). VPs exhibit a very broad substrate scope, and accordingly, we chose five substrates that represent different redox potentials and chemical structures: ABTS and 2,6-dimethoxyphenol (DMP; low-redox potential substrates), veratryl alcohol (VA; a high-redox potential substrate), reactive black 5 (RB5; a high-redox potential dye) and Mn^2+^. DMP, VA, and Mn^2+^ are likely to be native substrates of white-rot VPs, since VPs functionalize them to serve as mediators for lignin oxidation. VA and RB5 are exclusively oxidized by the high-redox potential surface-active tryptophan, while DMP and ABTS are oxidized both in the heme pocket (low-) or by the tryptophan radical (high-efficiency site). Furthermore, Mn^2+^ is oxidized in a distinct active site. Thus, the five substrates probe the reactivity and selectivity of each of the three VP active sites. Remarkably, despite the structural complexity, the lack of co-factors in the model structures and the very large number of mutations in each design, the designs oxidize all of the substrates (Figure 3). Thus, the combination of *ab initio* modeling and PROSS design is effective even in complex, multifunctional enzymes such as VPs.

**Figure 3.**
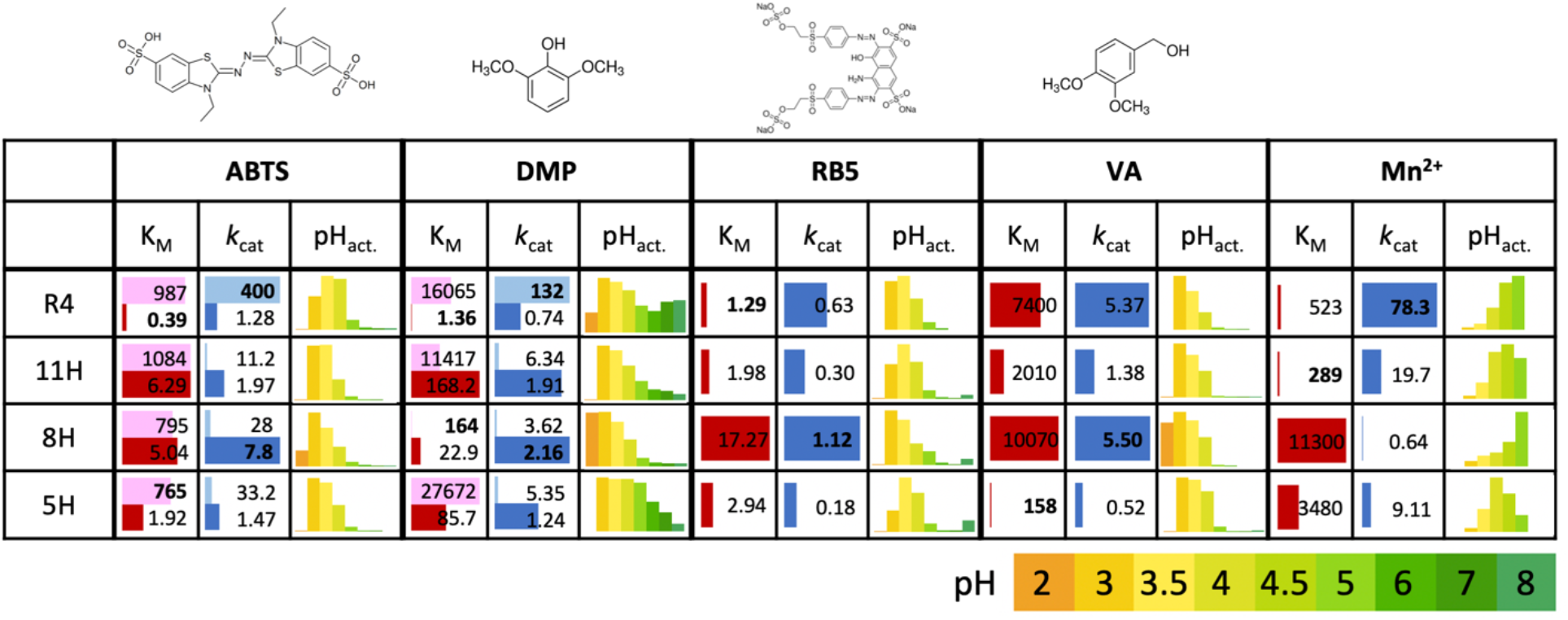
High functional diversity among VP designs. For each VP (5H, 8H, 11H and R4) and substrate (ABTS, DMP, RB5, VA and Mn^2+^), two kinetic parameters are shown, K_M_ and *k*_cat_ (in μM and sec^−1^, respectively); bars are normalized to the larger value in each substrate (worst K_M_ and best *k*_cat_). For ABTS and DMP, light bars refer to the kinetic parameters of the low-efficiency site and the dark bars to the high-efficiency site; kinetic values are normalized separately for each (Table S4 shows all kinetic parameters). The best affinity (lowest K_M_) and turnover number (highest *k*_cat_) for each substrate are highlighted in bold. pH-dependent activity profiles are shown for each enzyme-substrate pair, and the bars are normalized to the activity at optimal pH for each such pair (Figure S5 shows full activity plots).

The four VPs show the expected preference for acidic conditions (Figure 3 and Figure S5). Nevertheless, pH preferences vary dramatically among the designs. For instance, 8H exhibits a strong preference for low pH (2-3) in DMP and VA oxidation, whereas 5H is non-reactive at low pH and is reactive at relatively high pH where 8H is inactive. Furthermore, whereas R4 exhibits relatively broad pH reactivity for DMP, it is more restricted in pH scope on the other substrates relative to the designs. Manganese oxidation as a function of pH shows unexpected trends. Typically, VPs exhibit a pH optimum of 5 for oxidizing Mn^2+^ (as in R4)^11,13,26,30^. 8H also exhibits a pH optimum of 5, despite the fact that this design exhibits a more acidic pH optimum for the other substrates. By contrast, 5H and 11H have a pH optimum at 4 and 4.5, respectively, closer to the native conditions in which white-rot fungi operate (pH approximately 3^8^).

We next tested the reactivity profiles of the three designs and R4 relative to the substrates discussed above and hydrogen peroxide, again noting dramatic differences (Figure 3 and Table S4). 8H and R4 are the most efficient enzymes in terms of their turnover numbers (*k*_cat_), and R4 affinity to some substrates is the greatest, as reflected in low K_M_. Substrate affinities, however, exhibit surprising diversity without establishing any of the enzymes as best across the board. First, although 5H exhibits low *k*_cat_ values throughout, it exhibits more than an order of magnitude higher affinity for VA than the next best enzyme. Considering that VA has the highest redox potential of all substrates and is a non-phenolic component of lignin, the enzyme’s high affinity for VA may be valuable for complete decomposition of lignin at low VA concentrations. Second, 11H exhibits the highest affinity to H_2_O_2_ (by an order of magnitude relative to R4). Since H_2_O_2_ deactivates VPs and destabilizes proteins generally, high affinity in this enzyme is beneficial in cases where hydrogen peroxide levels must be kept low to maintain activity in other enzymes. Third, 11H exhibits the highest affinity for manganese, by at least twofold (relative to R4). Last, the heme-dependent DMP affinity of 8H is at least 70-fold higher than that of the other VPs.

### The structural basis for functional diversity in designed VPs

8H was designed based on a VP from the Polyporales order, whereas the other designs, as well as R4, derive from the Agaricales order (and specifically, from genus *Pleurotus*). Thus, 8H diverges from the other VPs both in sequence (Tables S2 & S3) and in structure (Figure 4A). The most significant active-site differences between 8H and the other VPs are in a loop that chelates both heme and manganese (Figure 4B). Among the manganese-chelating residues, 8H exhibits an Asp→His mutation in position 175 (all position numbers relative to PDB entry 3FJW), which was previously implicated in manganese oxidation^26^. Additionally, mutation Ala173Ser near the manganese binding site may also modify manganese-binding properties. These naturally occurring sequence changes likely explain the sharp decrease in manganese oxidation ability in 8H.

**Figure 4.**
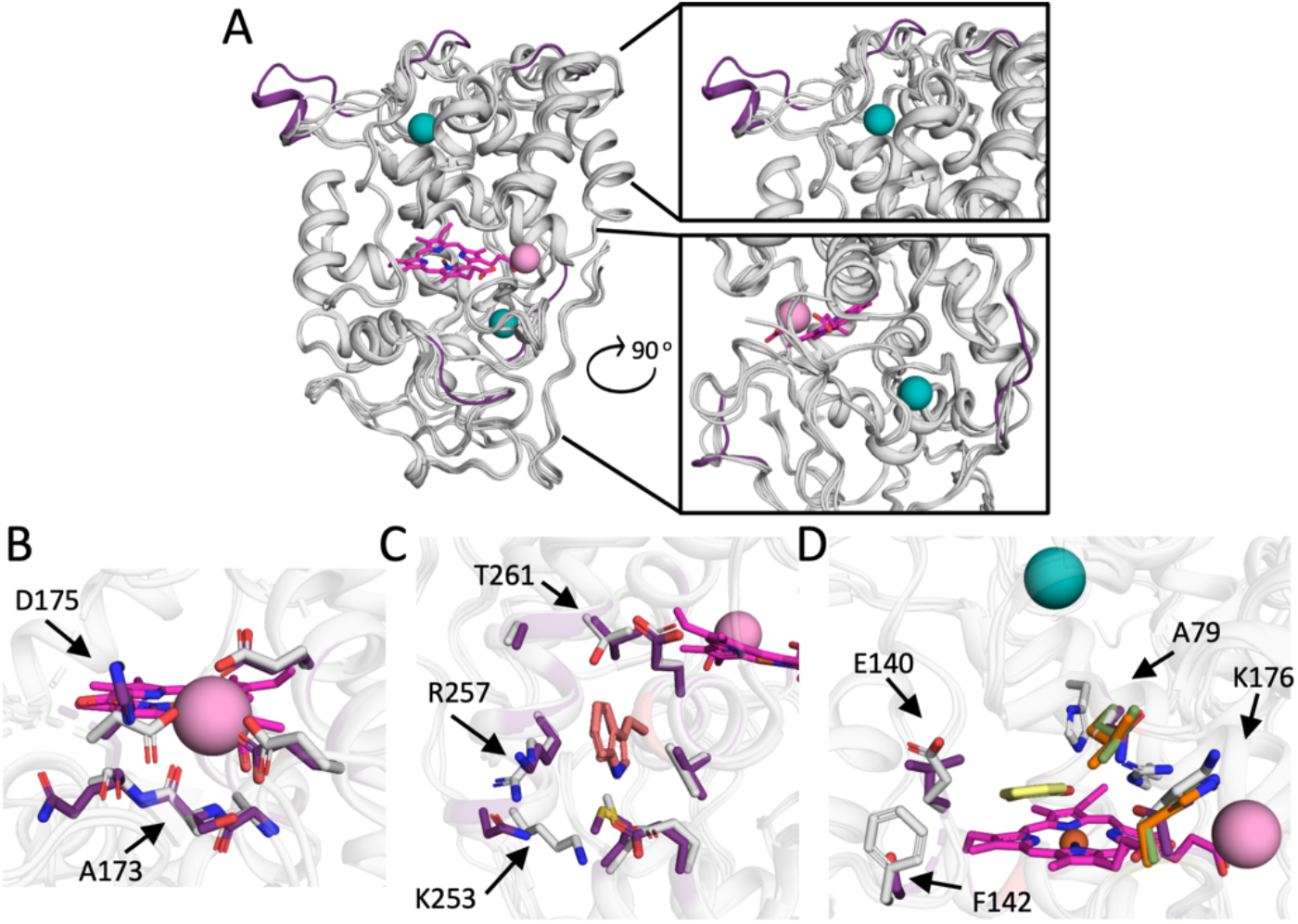
Structural basis for the different activity profiles in VPs. (A) AlphaFold2^22^ models of 5H, 8H and 11H are superimposed onto the VPL crystallographic structure (PDB entry 3FJW). All VP backbones are presented in gray cartoons, VPL calcium and manganese ions are in teal and pink spheres, respectively, and the heme group in pink sticks in all panels. Backbone segments that are unique for 8H are colored in purple. (B) Manganese-oxidation site. VPL and 8H residues that chelate manganese and vicinal side chains are in gray and purple sticks, respectively. Significant variations are marked in arrows and their position identities and numbers are relative to PDB entry 3FJW in all panels. (C) Reactive surface tryptophanyl site. Tryptophan is presented in salmon sticks. VPL and 8H residues in the tryptophan vicinity are presented in gray and purple sticks, respectively. (D) Access channel to the heme-oxidation site. Guaiacol (GUA), which is chemically similar to DMP, from a VPL crystallographic structure (PDB entry 4G05^31^) is in yellow sticks. VPL residues in the GUA vicinity are gray. Mutations relative to VPL are indicated in green, purple and orange sticks for 5H, 8H and 11H, respectively, and are marked by arrows.

In addition to the manganese oxidation site, VPs oxidize substrates through a high-redox surface tryptophan site via a long-range electron-transfer mechanism, and a low-redox potential heme-dependent site. The four VPs we investigated exhibit large variations in kinetic constants in the high-redox surface site. Thus, despite the high accessibility of this site, the tryptophan enviroment’s chemical properties and molecular recognition play an important role in determining reactivity at this site^9^. 8H shows unique electrostatic and structural properties at this site as well: whereas the other enzymes exhibit a conserved Lys and Arg that partially shield the tryptophan and a Thr residue on the opposite side, 8H has an Asn, Lys and Val, respectively (Figure 4C). Although 5H, 11H and R4 share an identical sequence at the high-redox potential tryptophan active site, they exhibit significant changes in reactivity. These reactivity differences may stem from sequence changes along the electron-relay path that connects the tryptophan radical and the heme-dependent site in ways that remain to be clarified.

Finally, the models show that the heme pocket is conserved among all the VPs. The only variations in its immediate surroundings are the manganese-bridging loop in 8H, a Ser→Ala substitution in position 168 in 5H and hydrophobic-to-hydrophobic substitutions at positions 152, 234 and 262 in all VPs. Given these relatively minor sequence changes, we speculate that the ABTS and DMP activity differences stem from changes in substrate accessibility to the heme-binding pocket. Indeed, in heme access-channel loops, dramatic variations exist in several positions which influence the hydrophobicity of the pocket and its size. For example, 8H has a Leu instead of Glu in position 140, generating a hydrophobic access channel that may underlie the observed higher affinity for DMP; this mutation in 8H may also lead to higher affinity for the product, explaining the low turnover number (Figure 4D). Previous mutation analyses demonstrated this position’s importance for substrate recognition and pH-dependent activity^14,31^. We conclude that the significant diversity observed in the active sites and access channels among natural VPs underlies the large functional changes observed among our designs. These structure-activity relationships may inform future VP design efforts or help focus the search for natural VPs that exhibit unique functional features.

## Discussion

Lignocellulose is a heterogenous and stable material. To effectively decompose it, white-rot fungi deploy an arsenal of oxidoreductases that act synergistically, including through consortia comprising different fungal species^8,10,32,33^. One of the major research goals in mobilizing lignocellulose for energy production is to deploy such an arsenal in a heterologous or cell-free setting^34^. In the case of VPs, however, this goal is stymied by the lack of a common host for functional expression, thus demanding extensive enzyme engineering. Current enzyme-engineering approaches, however, are impossible to apply in cases in which the wild type protein exhibits no functional expression in commonly used heterologous hosts as is the case for the VPs we targeted. Accordingly, VP engineering studies have focused on VPL and used directed evolution to isolate variants that exhibited higher expressibility, stability and pH-dependent activity^13–15^. We demonstrated that deep-learning *ab initio* structure prediction methods can be combined seamlessly with the PROSS protein-stability design method to deliver new VPs that add significant diversity to the arsenal of lignin oxidoreductases that have been characterized to date. Our structural analysis suggests that the observed functional diversity stems from the natural sequence diversity and that the design process serves only to expose and utilize this preexisting diversity.

The designs exhibit a number of attractive features for use in a synergistic oxidoreductase cocktail. First, they are thermostable and relatively well expressed in *S. cerevisiae*. Second, they exhibit different pH tolerance and pH-dependent activity profiles that can be useful in different research or applied settings. Third, they show dramatic differences in substrate selectivity. Even with respect to a given substrate, the enzymes exhibit large differences in catalytic turnover and substrate affinity (*k*_cat_ and k_M_, respectively). By combining enzymes with high and low ligand affinity (and low and high turnover, respectively), one may generate a cocktail that can oxidize substrates quickly when substrate concentrations are high (through the high-turnover enzyme) and continue to oxidize substrates even when substrate concentrations are limiting (through the high-affinity enzyme), thus maximizing the cocktail’s effectiveness^35,36^.

Finally, we note the remarkably high accuracy of the new generation of deep-learning based structure predictors. Prior to using the Dec. 2020 version of trRosetta, we attempted modeling the VPs using a previous version of trRosetta and also using traditional homology-modelling software. The resulting structure models, however, did not exhibit the features expected of a VP enzyme and we did not pursue them further. The most successful designs in our study comprised approximately 40 mutations from the wild type proteins, underscoring the high accuracy and reliability of the structure prediction and design methods we combined. Recently, even more reliable predictors have been made publicly available^22,23^. Our work demonstrates that enzyme engineering can now be freed from the requirement of experimental structure determination through a streamlined pipeline that starts from *ab initio* modeling and continues to automated protein design. Accordingly, we have adapted the PROSS web server (https://pross.weizmann.ac.il/) to use AlphaFold2 models, automatically restricting design to high-confidence regions in the models. Compared to previous VP engineering studies that required screening thousands of variants^13–15^, we screened fewer than 40 designs. Thus, model-based enzyme design may address the growing need in many fields to generate functional, stable, and efficient enzyme repertoires or pathways in one shot^37^.

## Acknowledgments

We thank members of the Fleishman lab for comments and advice. Research in the Fleishman lab was supported by an Individual Grant from the Israel Science Foundation (1844/19), a Consolidator Award from the European Research Council (815379), by the Sustainability and Energy Research Initiative, and by a donation in memory of Sam Switzer. SB-Z and SJF are named inventors on a patent filing describing the VP designs, and SJF is a named inventor on patents describing PROSS and various PROSS designs.

## References

(1) Ragauskas, A. J.; Beckham, G. T.; Biddy, M. J.; Chandra, R.; Chen, F.; Davis, M. F.; Davison, B. H.; Dixon, R. A.; Gilna, P.; Keller, M.; Langan, P.; Naskar, A. K.; Saddler, J. N.; Tschaplinski, T. J.; Tuskan, G. A.; Wyman, C. E. Lignin Valorization: Improving Lignin Processing in the Biorefinery. Science 2014, 344 (6185), 1246843.

(2) Ragauskas, A. J.; Williams, C. K.; Davison, B. H.; Britovsek, G.; Cairney, J.; Eckert, C. A.; Frederick, W. J., Jr; Hallett, J. P.; Leak, D. J.; Liotta, C. L.; Mielenz, J. R.; Murphy, R.; Templer, R.; Tschaplinski, T. The Path Forward for Biofuels and Biomaterials. Science 2006, 311 (5760), 484–489.

(3) Tuck, C. O.; Pérez, E.; Horváth, I. T.; Sheldon, R. A.; Poliakoff, M. Valorization of Biomass: Deriving More Value from Waste. Science 2012, 337 (6095), 695–699.

(4) Davidi, L.; Moraïs, S.; Artzi, L.; Knop, D.; Hadar, Y.; Arfi, Y.; Bayer, E. A. Toward Combined Delignification and Saccharification of Wheat Straw by a Laccase-Containing Designer Cellulosome. Proc. Natl. Acad. Sci. U. S. A. 2016, 113 (39), 10854–10859.

(5) da Costa Sousa, L.; Chundawat, S. P. S.; Balan, V.; Dale, B. E. “Cradle-to-Grave” Assessment of Existing Lignocellulose Pretreatment Technologies. Curr. Opin. Biotechnol. 2009, 20 (3), 339–347.

(6) Chen, C.-C.; Dai, L.; Ma, L.; Guo, R.-T. Enzymatic Degradation of Plant Biomass and Synthetic Polymers. Nature Reviews Chemistry 2020, 4 (3), 114–126.

(7) Martínez, A. T.; Ruiz-Dueñas, F. J.; Martínez, M. J.; Del Río, J. C.; Gutiérrez, A. Enzymatic Delignification of Plant Cell Wall: From Nature to Mill. Curr. Opin. Biotechnol. 2009, 20 (3), 348–357.

(8) Alcalde, M. Engineering the Ligninolytic Enzyme Consortium. Trends Biotechnol. 2015, 33 (3), 155–162.

(9) Ruiz-Dueñas, F. J.; Morales, M.; García, E.; Miki, Y.; Martínez, M. J.; Martínez, A. T. Substrate Oxidation Sites in Versatile Peroxidase and Other Basidiomycete Peroxidases. J. Exp. Bot. 2009, 60 (2), 441–452.

(10) Ruiz-Dueñas, F. J.; Lundell, T.; Floudas, D.; Nagy, L. G.; Barrasa, J. M.; Hibbett, D. S.; Martínez, A. T. Lignin-Degrading Peroxidases in Polyporales: An Evolutionary Survey Based on 10 Sequenced Genomes. Mycologia 2013, 105 (6), 1428–1444.

(11) Ayuso-Fernández, I.; Ruiz-Dueñas, F. J.; Martínez, A. T. Evolutionary Convergence in Lignin-Degrading Enzymes. Proc. Natl. Acad. Sci. U. S. A. 2018, 115 (25), 6428–6433.

(12) Pérez-Boada, M.; Ruiz-Dueñas, F. J.; Pogni, R.; Basosi, R.; Choinowski, T.; Martínez, M. J.; Piontek, K.; Martínez, A. T. Versatile Peroxidase Oxidation of High Redox Potential Aromatic Compounds: Site-Directed Mutagenesis, Spectroscopic and Crystallographic Investigation of Three Long-Range Electron Transfer Pathways. J. Mol. Biol. 2005, 354 (2), 385–402.

(13) Garcia-Ruiz, E.; Gonzalez-Perez, D.; Ruiz-Dueñas, F. J.; Martínez, A. T.; Alcalde, M. Directed Evolution of a Temperature-, Peroxide- and Alkaline pH-Tolerant Versatile Peroxidase. Biochem. J 2012, 441 (1), 487–498.

(14) Gonzalez-Perez, D.; Mateljak, I.; Garcia-Ruiz, E.; Ruiz-Dueñas, F. J.; Martinez, A. T.; Alcalde, M. Alkaline Versatile Peroxidase by Directed Evolution. Catal. Sci. Technol. 2016, 6 (17), 6625–6636.

(15) Gonzalez-Perez, D.; Garcia-Ruiz, E.; Ruiz-Dueñas, F. J.; Martinez, A. T.; Alcalde, M. Structural Determinants of Oxidative Stabilization in an Evolved Versatile Peroxidase. ACS Catal. 2014, 4 (11), 3891–3901.

(16) Weinstein, J.; Khersonsky, O.; Fleishman, S. J. Practically Useful Protein-Design Methods Combining Phylogenetic and Atomistic Calculations. Curr. Opin. Struct. Biol. 2020, 63, 58–64.

(17) Goldenzweig, A.; Goldsmith, M.; Hill, S. E.; Gertman, O.; Laurino, P.; Ashani, Y.; Dym, O.; Unger, T.; Albeck, S.; Prilusky, J.; Lieberman, R. L.; Aharoni, A.; Silman, I.; Sussman, J. L.; Tawfik, D. S.; Fleishman, S. J. Automated Structure- and Sequence-Based Design of Proteins for High Bacterial Expression and Stability. Mol. Cell 2016, 63 (2), 337–346.

(18) Weinstein, J. J.; Goldenzweig, A.; Hoch, S.-Y.; Fleishman, S. J. PROSS 2: A New Server for the Design of Stable and Highly Expressed Protein Variants. Bioinformatics 2020. https://doi.org/10.1093/bioinformatics/btaa1071.

(19) Peleg, Y.; Vincentelli, R.; Collins, B. M.; Chen, K.-E.; Livingstone, E. K.; Weeratunga, S.; Leneva, N.; Guo, Q.; Remans, K.; Perez, K.; Bjerga, G. E. K.; Larsen, Ø.; Vaněk, O.; Skořepa, O.; Jacquemin, S.; Poterszman, A.; Kjær, S.; Christodoulou, E.; Albeck, S.; Dym, O.; Ainbinder, E.; Unger, T.; Schuetz, A.; Matthes, S.; Bader, M.; de Marco, A.; Storici, P.; Semrau, M. S.; Stolt-Bergner, P.; Aigner, C.; Suppmann, S.; Goldenzweig, A.; Fleishman, S. J. Community-Wide Experimental Evaluation of the PROSS Stability-Design Method. J. Mol. Biol. 2021, 433 (13), 166964.

(20) Zahradník, J.; Kolářová, L.; Peleg, Y.; Kolenko, P.; Svidenská, S.; Charnavets, T.; Unger, T.; Sussman, J. L.; Schneider, B. Flexible Regions Govern Promiscuous Binding of IL-24 to Receptors IL-20R1 and IL-22R1. FEBS J. 2019. https://doi.org/10.1111/febs.14945.

(21) Mao, G.; Wang, F.; Wang, J.; Chen, P.; Zhang, X.; Zhang, H.; Wang, Z.; Song, A. A Sustainable Approach for Degradation and Detoxification of Malachite Green by an Engineered Polyphenol Oxidase at High Temperature. J. Clean. Prod. 2021, 328, 129437.

(22) Jumper, J.; Evans, R.; Pritzel, A.; Green, T.; Figurnov, M.; Ronneberger, O.; Tunyasuvunakool, K.; Bates, R.; Žídek, A.; Potapenko, A.; Bridgland, A.; Meyer, C.; Kohl, S. A. A.; Ballard, A. J.; Cowie, A.; Romera-Paredes, B.; Nikolov, S.; Jain, R.; Adler, J.; Back, T.; Petersen, S.; Reiman, D.; Clancy, E.; Zielinski, M.; Steinegger, M.; Pacholska, M.; Berghammer, T.; Bodenstein, S.; Silver, D.; Vinyals, O.; Senior, A. W.; Kavukcuoglu, K.; Kohli, P.; Hassabis, D. Highly Accurate Protein Structure Prediction with AlphaFold. Nature 2021, 596 (7873), 583–589.

(23) Baek, M.; DiMaio, F.; Anishchenko, I.; Dauparas, J.; Ovchinnikov, S.; Lee, G. R.; Wang, J.; Cong, Q.; Kinch, L. N.; Schaeffer, R. D.; Millán, C.; Park, H.; Adams, C.; Glassman, C. R.; DeGiovanni, A.; Pereira, J. H.; Rodrigues, A. V.; van Dijk, A. A.; Ebrecht, A. C.; Opperman, D. J.; Sagmeister, T.; Buhlheller, C.; Pavkov-Keller, T.; Rathinaswamy, M. K.; Dalwadi, U.; Yip, C. K.; Burke, J. E.; Garcia, K. C.; Grishin, N. V.; Adams, P. D.; Read, R. J.; Baker, D. Accurate Prediction of Protein Structures and Interactions Using a Three-Track Neural Network. Science 2021, 373 (6557), 871–876.

(24) Yang, J.; Anishchenko, I.; Park, H.; Peng, Z.; Ovchinnikov, S.; Baker, D. Improved Protein Structure Prediction Using Predicted Interresidue Orientations. Proc. Natl. Acad. Sci. U. S. A. 2020, 117 (3), 1496–1503.

(25) Hiranuma, N.; Park, H.; Baek, M.; Anishchenko, I.; Dauparas, J.; Baker, D. Improved Protein Structure Refinement Guided by Deep Learning Based Accuracy Estimation. Nat. Commun. 2021, 12 (1), 1340.

(26) Ruiz-Dueñas, F. J.; Morales, M.; Pérez-Boada, M.; Choinowski, T.; Martínez, M. J.; Piontek, K.; Martínez, A. T. Manganese Oxidation Site in Pleurotus Eryngii Versatile Peroxidase: A Site-Directed Mutagenesis, Kinetic, and Crystallographic Study. Biochemistry 2007, 46 (1), 66–77.

(27) Lú-Chau, T. A.; Ruiz-Dueñas, F. J.; Camarero, S.; Feijoo, G.; Martínez, M. J.; Lema, J. M.; Martínez, A. T. Effect of pH on the Stability of Pleurotus Eryngii Versatile Peroxidase during Heterologous Production in Emericella Nidulans. Bioprocess Biosyst. Eng. 2004, 26 (5), 287–293.

(28) Gao, Y.; Zheng, L.; Li, J.-J.; Du, Y. Insight into the Impact of Two Structural Calcium Ions on the Properties of Pleurotus Eryngii Versatile Ligninolytic Peroxidase. Arch. Biochem. Biophys. 2016, 612, 9–16.

(29) Valderrama, B.; Ayala, M.; Vazquez-Duhalt, R. Suicide Inactivation of Peroxidases and the Challenge of Engineering More Robust Enzymes. Chem. Biol. 2002, 9 (5), 555–565.

(30) Knop, D.; Levinson, D.; Makovitzki, A.; Agami, A.; Lerer, E.; Mimran, A.; Yarden, O.; Hadar, Y. Limits of Versatility of Versatile Peroxidase. Appl. Environ. Microbiol. 2016, 82 (14), 4070–4080.

(31) Morales, M.; Mate, M. J.; Romero, A.; Martínez, M. J.; Martínez, Á. T.; Ruiz-Dueñas, F. J. Two Oxidation Sites for Low Redox Potential Substrates: A Directed Mutagenesis, Kinetic, and Crystallographic Study on Pleurotus Eryngii Versatile Peroxidase. J. Biol. Chem. 2012, 287 (49), 41053–41067.

(32) Sethupathy, S.; Morales, G. M.; Li, Y.; Wang, Y.; Jiang, J.; Sun, J.; Zhu, D. Harnessing Microbial Wealth for Lignocellulose Biomass Valorization through Secretomics: A Review. Biotechnol. Biofuels 2021, 14 (1), 154.

(33) Wang, J.; Li, L.; Xu, H.; Zhang, Y.; Liu, Y.; Zhang, F.; Shen, G.; Yan, L.; Wang, W.; Tang, H.; Qiu, H.; Gu, J.-D.; Wang, W. Construction of a Fungal Consortium for Effective Degradation of Rice Straw Lignin and Potential Application in Bio-Pulping. Bioresour. Technol. 2021, 344 (Pt B), 126168.

(34) Raulo, R.; Heuson, E.; Froidevaux, R.; Phalip, V. Combining Analytical Approaches for Better Lignocellulosic Biomass Degradation: A Way of Improving Fungal Enzymatic Cocktails? Biotechnol. Lett. 2021. https://doi.org/10.1007/s10529-021-03201-2.

(35) Eisenthal, R.; Danson, M. J.; Hough, D. W. Catalytic Efficiency and kcat/KM: A Useful Comparator? Trends Biotechnol. 2007, 25 (6), 247–249.

(36) Carrillo, N.; Ceccarelli, E.; Roveri, O. Usefulness of Kinetic Enzyme Parameters in Biotechnological Practice. Biotechnol. Genet. Eng. Rev. 2010, 27, 367–382.

(37) Ospina, F.; Schülke, K. H.; Hammer, S. C. Biocatalytic Alkylation Chemistry: Building Molecular Complexity with High Selectivity. ChemPlusChem 2021. https://doi.org/10.1002/cplu.202100454.

