## Supplemental Information for "Stable and functionally diverse versatile peroxidases by computational design directly from sequence"

#### Methods

**VP sequence collection.** VP sequences were extracted from three databases: MycoCosm (fungal genome database[1]), RedoxiBase (oxidoreductases database[2]), and fPoxDB (fungal peroxidases database[3]). Signal peptide sequences were identified using SignalP-5.0[4] and were removed from the relevant sequences. A multiple sequence alignment was generated using MUSCLE[5] based on all unique sequences, and a phylogenetic tree was inferred using the Maximum Likelihood method and JTT matrix-based model through MEGA[6,7]. For each clade, a consensus sequence was generated using EMBOSS Cons[8], and the VP sequence with the highest similarity to the consensus was chosen for further analysis.

**trRosetta structure modeling.** The structures of all the selected sequences were calculated using the trRosetta structure prediction algorithm (December 2020 version[9,10]) through the Robetta server (https://robetta.bakerlab.org/). The models were visually inspected, and ones exhibiting poor parameters were eliminated; for instance, models that did not adopt the expected fold of VPs or in which the catalytic site diverged from expectation. Finally, twelve diverse sequences (eleven sequences from the analysis above and VPL), with 51-81% identity between each sequence pair, were selected for further modeling and design (Table S1 & S2).

**PROSS stability design calculations.** For each sequence, the best-calculated model was subjected to design by the PROSS algorithm[11,12]. VPL was also designed by PROSS starting from its crystal structure (PDB entry: 3FJW). Since trRosetta does not predict ligands or ions, structural information from the VPL structure was used to determine the putative binding sites of the heme, manganese, and two structural calcium ions, as well as of the active tryptophan, and to limit mutations in these sites. Design in positions that showed structural inconsistency in the best five models calculated by trRosetta and in positions that are in contact with such positions, was disallowed as well. Automated filtering of such positions based on AlphaFold2 models is now enabled on the PROSS web server (https://PROSS.weizamann.ac.il). Further, any mutations that might change native N-glycosylation patterns were dismissed.

**AlphaFold2 modelling.** The sequences of 5H, 8H and 11H were structurally modeled by the AlphaFold2 algorithm[13], using the AlphaFold Colab notebook:
https://colab.research.google.com/github/deepmind/alphafold/blob/main/notebooks/AlphaFold.ipynb

**Reagents.** The protease-deficient *S. cerevisiae* strain BJ5465 (α ura3-52 trp1 leu2Δ1 his3Δ200 pep4::HIS3 prb1Δ1.6R can1 GAL) was obtained from LGCPromochem (Barcelona, Spain). The uracil-independent and ampicillin resistance shuttle vector pJRoC30 was obtained from the California Institute of Technology (Caltech, Pasadena, CA). The α-factor prepro-leader sequence and all the VP genes sequences were ordered from Twist Biosciences (San Francisco, CA). The BamHI and XhoI restriction enzymes were ordered from New England Biolabs (NEB, Rehovot, Israel). ABTS, VA, RB5, and the *S. cerevisiae* transformation kit, were purchased from Sigma–Aldrich (Rehovot, Israel). DMP was purchased from Acros Organic (Geel, Belgium) and hemoglobin from bovine erythrocytes from EMD Millipore Corp (Billerica, MA, USA).

**Cloning of VP genes.** Cloning of all VP genes was performed by using the *S. cerevisiae* homologous recombination machinery[14]. pJRoC30-AAO (aryl-alcohol oxidase) expression shuttle vector previously constructed in the Alcalde lab[15] was digested with BamHI and XhoI restriction enzymes to remove the signal peptide and the AAO gene constructed within it. The α-factor prepro-leader DNA sequence, that was used in a previous directed evolution campaign of VPL (including additional restriction site in its 3' that encodes for Glu-Phe dipeptide in the N-terminal of the mature proteins)[16], was ordered as a gene fragment with 40 bp overlap to the linearized plasmid, and the VP genes were ordered each with 40 bp overlap to the signal peptide sequence, and to the linearized plasmid. The design of the 40 bp overlapping regions between the three fragments (plasmid, signal peptide, VP genes) allowed the recombination machinery of the protease-deficient *S. cerevisiae* strain BJ5465 to drive the fusion of the three DNA elements after transformation, and to form the pJRoC30-SignalPeptide-VPgene expression shuttle vector. pJRoC30-VPL-WT, -R4, and -2-1B were constructed previously in the Alcalde lab[16]. All *S. cerevisiae*-transformed cells were plated in synthetic complete (SC) drop-out plates, and in each plate, selected colonies were picked and sequenced to verify the correct assembly and gene sequence.

**Culture media.** Minimal medium is composed of 6.7 g/L yeast nitrogen base, 1.92 g/L amino acids supplements (yeast synthetic drop-out medium supplements without uracil), 2% raffinose and 25 mg/L chloramphenicol. VPs expression medium is composed of YP x1.11 medium (22.2 g/L bacto peptone and 11.1 g/L yeast extract), 67 mM $KH_2PO_4$ buffer at pH 6.0, 25 g/L ethanol, 22.2 g/L D-galactose, 500 mg/L bovine hemoglobin, 1 mM $CaCl_2$ and 25 mg/L chloramphenicol. SC drop-out plates are composed of 6.7 g/L yeast nitrogen base, 1.92 g/L amino acids supplements (yeast synthetic drop-out medium supplements without uracil), 2% glucose, 20 g/L Bacto agar and 25 mg/L chloramphenicol.

**Screening for active variants.** A colony from each *S. cerevisiae* clone containing the parental or mutant VP gene was picked from an SC drop-out plate, inoculated in 2 mL minimal medium in a 14 mL culture tube, and incubated for 48 hours at 30 °C and 225 rpm. An aliquot of cells was removed and used to inoculate 2 mL of minimal medium in a new 14 mL culture tube to an $OD_{600nm}$ of 0.25-0.30, under the same conditions. The cells completed two growth phases (8-10 hours, reaching $OD_{600nm} \sim 1$), then the expression medium (2.7 mL) was inoculated with 0.3 mL of the pre-culture in a new 14 mL culture tube ($OD_{600nm} \sim 0.1$). Cells were incubated for further ~38-40 hours at 30 °C and 225 rpm and then centrifuged at 4000 g for 20 min at 4°C. The supernatant was removed into new tubes for further analysis. The expression protocol ran in triplicate, with an empty vector (containing only the signal peptide sequence) and the VPL-R4 and -2-1B variants as negative and positive controls, respectively. An ABTS-based colorimetric assay was conducted to assess the variants' activity: 20 µL of supernatant were transferred into activity 96 plates (Greiner Bio-One GmbH, Kremsmünster, Austria), and then, 180 µL of the reaction mixture were added to each row in the plate, and absorption at 418 nm was recorded immediately in a kinetic mode in a plate-reader at 25 °C (Citation5 or Synergy HTX plate readers, Bio-Tek, Bad Friedrichshall, Germany). The reaction mixture contained 100 mM citrate–phosphate buffer (pH 4.0), 2 mM ABTS and 0.1 mM $H_2O_2$. The activities were recorded in triplicate.

**Small scale production of active variants.** A colony from each *S. cerevisiae* clone containing the parental or mutant VP gene was picked from an SC drop-out plate, inoculated in 2 mL minimal medium in a 14 mL culture tube, and incubated for 48 hours at 30 °C and 225 rpm. An aliquot of cells was removed and used to inoculate 2.7 mL of minimal medium in a new 14 mL culture tube to an $OD_{600nm}$ of 0.25-0.30, under the same conditions. The cells completed two growth phases (8-10 hours, reaching $OD_{600nm}$ ~ 1), then the expression medium (9 mL) was inoculated with 1 mL of the pre-culture in a 50 mL Falcon tube ($OD_{600nm}$ ~ 0.1). Cells were incubated for further ~60 hours at 30 °C and 225 rpm and then centrifuged at 4000 g for 20 min at 4°C. The supernatant was removed into new tubes for further characterization of VPs in supernatant.

**Thermostability assay ($T_{50}$).** Aliquots of 30 µL of selected variants' supernatant at appropriate dilutions (with 20 mM piperazine pH=5.5, buffer A, to achieve linear response in kinetic mode measurements in activity reads) were used for each incubation temperature. The samples were incubated for 10 or 15 minutes in a thermocycler pre-heated to a specific temperature (every 5 °C in a gradient scale ranging from 25 to 80 °C) and then removed and chilled on ice for 10 min. Thereafter, samples were removed from ice and incubated for at least 5 min at room temperature. Activity at each temperature was measured using the ABTS-based colorimetric assay described above and was normalized to the activity at 25 °C for residual activity calculations. All incubations and activity assays were conducted in triplicate. $T_{50}$ values were calculated by sigmoidal fit to the $T_{50}$ data of 5H, 8H, 11H and R4 (15 minutes incubation).

**Kinetic thermostability ($t_{1/2}$).** Aliquots of 30 µL of selected variants' supernatant at appropriate dilutions (with buffer A, to achieve linear response in kinetic mode measurements in activity reads) were used for each incubation time point. The samples were incubated in a thermocycler (S1000$^{TM}$ thermocycler, Bio-Rad, Rishon LeZion, Israel) pre-heated to 60 °C or 65 °C, and removed at different times (after 0, 2, 5, 7, 10, 15, 20, 30, 45, 60, 90 and 120 min), chilled out on ice for 10 min and further incubated at room temperature at least for 10 min. Activity at each time point was measured using the ABTS-based colorimetric assay described above and was normalized to the activity at time 0 for residual activity calculations. All incubations and activity assays were conducted in triplicate.

**pH stability.** Supernatants of the selected variants were diluted to reach a final concentration of 100 mM citrate-phosphate-borate buffer at pH ranging from 2-9. Aliquots of 20 µL were removed at different times (time 0, 4, 25, 50, 75 and 165 hours) and measured in the regular ABTS-based colorimetric assay described above, but here in the presence of 180 µL of the following reaction mixture: 111.11 mM citrate-phosphate-borate buffer (pH 4.0), 2.22 mM ABTS and 0.111 mM $H_2O_2$. For pH 2, an additional experiment was conducted, under the same procedure but with aliquots being removed at different time points (time 0, 10, 20, 30, 45, 60, 75, 90, 120, and 150 min). For the assay in pH range 2-9, activities were normalized to the activity at time 0 in pH=3, for residual activity calculations. All incubations and activity assays were conducted in triplicate.

**VPs production and purification.** A colony from *S. cerevisiae* clone containing the VP gene (5H, 8H, 11H, VPL_R4, VPL_2-1B) was picked from an SC drop-out plate, inoculated in 25 mL minimal medium in a 250 flask, and incubated for 48 hours at 30 °C and 225 rpm. An aliquot of cells was removed and used to inoculate 100 mL of minimal medium in a 1 L flask to an $OD_{600nm}$ of 0.25-0.30, under the same conditions. The cells completed two growth phases (8-10 hours, reaching $OD_{600nm}$ ~ 1-1.5), then the expression medium (450 mL) was inoculated with 50 mL of the pre-culture in a 2 L flask ($OD_{600nm}$ ~ 0.1). Cells were incubated for a further ~60 hours at 30 °C and 225 rpm. Thereafter, cells were centrifuged at 6000 g for 15 min at 4°C, and the supernatant was collected and filtered with a 0.2 µm filter bottle.

Filtrates were subjected to fractional precipitation with ammonium sulfate in two steps: a first cut of 50%, followed by centrifugation and elimination of the precipitates, and a second cut of 70%. Buffer A was used to dissolve the pellet of the second cut, and the dissolved protein solution was shaken overnight at 4 $^{o}$C for maximal recovery. The dissolved fraction was then centrifuged, filtrated, concentrated, and subjected to overnight dialysis against buffer A. Filtered fractions of the VP proteins after dialysis were uploaded into a HiTrap$^{TM}$ Q HP Column (GE Healthcare Bio-Sciences AB, Uppsala, Sweden) pre-equilibrated with buffer A, through ÄKTA pure protein purification system (GE Healthcare Bio-Sciences AB). Proteins were eluted in a two-step linear gradient from 0 to 1 M NaCl, at a flow rate of 1 mL/min: the first phase of 0-25 % over 15 column volumes (75 min) and second phase of 25-100 % over 2 column volumes (10 min). The fractions of the peak with the highest VP activity (and absorption at 407 nm) were pooled, concentrated, and dialyzed against 20 mM piperazine buffer pH=5.5 and 150 mM NaCl (buffer B). Protein fractions were then uploaded onto a Superdex 75 Increase 10/300 GL (GE Healthcare Bio-Sciences AB) through the ÄKTA pure system pre-equilibrated with buffer B. The fractions of the peak with the highest VP activity (and absorption at 407 nm) were pooled and dialyzed against buffer A. Pure protein samples were stored at 4 °C. Protein concentration was determined using the BCA assay with bovine serum albumin as a standard. The obtained Reinheitszahl values (Rz: $Abs_{407\ nm}/Abs_{280\ nm}$), which indicate for the purity of peroxidases, were 1 for 8H (due to high tryptophan content and therefore extinction coefficient at 280 nm) and above 2 for 5H, 11H, R4 and 2-1B.

**Hydrogen peroxide stability.** Purified 5H, 8H, 11H, R4 and 2-1B at 250 nM concentration (diluted with buffer A, to achieve linear response in kinetic mode measurements in activity reads) were incubated for 50 minutes at room temperature with 750 µM $H_2O_2$ (1:3,000 molar ratio). An aliquot of 20 µL was removed at times 0, 3, 7, 12, 18, 28, 38 and 48 minutes. Activity was immediately measured using the ABTS-based colorimetric assay described above and was normalized to the activity at time zero for residual activity calculations. All incubations and activity assays were conducted in triplicate.

**pH activity profiles.** For purified 5H, 8H, 11H, and R4, 20 µL protein samples (diluted in buffer A) were transferred into activity 96 plates (in the case of VA and $MnSO_4$, UV-Star plates; Greiner Bio-One GmbH, Kremsmünster, Austria) and then, 180 µL of the reaction mixture were added to each row in the plate, and absorption at the appropriate wavelength (substrate-dependent) was recorded immediately in a kinetic mode in a plate-reader at 25 °C. The reaction mixtures contained a specific substrate in 100 mM citrate–phosphate-borate buffer (pH 2, 3, 3.5, 4, 5, 6, 7, 8) and 0.1 mM $H_2O_2$. The activities were recorded in triplicate. The following substrate concentrations and absorption wavelengths were used: ABTS: 2 mM, 418 nm; DMP: 5 mM, 469 nm; RB5: 0.05 mM, 598 nm; VA: 30 mM, 310 nm. For manganese, the reaction mixtures contained 60 mM $MnSO_4$ in 100 mM sodium tartrate buffer (pH 3, 3.5, 4, 4.5, 5) and 0.1 mM $H_2O_2$, and the $Mn^{3+}$-tartrate complex absorption was read at 238 nm. For each protein and substrate, the activities were normalized to the activity at optimal pH for residual activity calculations. Each activity assay was conducted in triplicate.

**Kinetic parameters.** Steady-state kinetics were determined for 5H, 8H, 11H and R4, by measuring the activity (initial rates) in increasing concentrations of the substrate, and the $K_M$ and $k_{cat}$ values were calculated by fitting the results to the Michaelis-Menten model ($V_0 = k_{cat}[E][S] / (K_M + [S])$). 20 µL purified protein samples (diluted in buffer A to appropriate concentration, [E]<<[S]) were transferred into activity 96 plates (in the case of VA and $MnSO_4$, UV-Star plates) and then, 180 µL of the reaction mixture were added to each row in the plate, and absorption at the appropriate wavelength (substrate-dependent) was recorded immediately in a kinetic mode in a plate-reader at 25 °C. The reaction mixtures contained substrates at varying concentrations, in 100 mM citrate–phosphate-borate buffer at optimum pH (for manganese, sodium tartrate buffer was used) and optimum $H_2O_2$ concentration (0.4 mM for 5H, 0.2 mM for 8H, 0.1 mM for 11H and 1 mM for R4; approximately double of the $K_M$ values were used to gain high activity with minimal inhibition effect). $H_2O_2$ kinetics was measured using 2 mM (5H, 11H and R4) or 3 mM (8H) ABTS in 100 mM citrate–phosphate-borate buffer at optimum pH for ABTS activity. The following molar extinction coefficients were used to calculate the substrate/product concentration: ABTS, $\varepsilon_{418\ nm} = 36{,}000\ M^{-1}\ cm^{-1}$; DMP, $\varepsilon_{469\ nm} = 27{,}500\ M^{-1}\ cm^{-1}$; RB5, $\varepsilon_{598\ nm} = 30{,}000\ M^{-1}\ cm^{-1}$; VA, $\varepsilon_{310\ nm} = 9300\ M^{-1}\ cm^{-1}$; $Mn^{3+}$-tartrate, $\varepsilon_{238\ nm} = 6500\ M^{-1}\ cm^{-1}$. All activities were recorded in triplicate and the average velocity was used for the kinetic constants calculations.

**Expression level calculations.** Enzyme concentrations, [E], were extracted from the Michaelis-Menten equation ($V_0 = k_{cat}[S][E] / (K_M + [S])$), where $V_0$ are the initial rates observed during the screening experiments (means of biological triplicates) and [S] is the ABTS concentration at this experiment. The kinetic constants correspond to the calculated $K_M$ and $k_{cat}$ for ABTS as reported in Table S4 (of the low-efficiency site that dominates the reaction at the used ABTS concentration). The activity assay was performed at pH=4.0, therefore the initial activities were normalized to the activity at optimal pH (using the data from pH-dependent activity assay; Figure S5).

**Amino acid sequences of characterized VP designs:**

5H:
VSLPQKRATCSGGQTTSNEACCVLFDLMEDLQKNLFDGGQCGEQAHEALRLTFHDAIGFSPSRG
VMGGADGSVITFSDIETNFPANLGIDDIVEAEKSFLQRHNISAGDLVHFAATLAVTNCPGAPRIPFF
LGRPPATAPSPPGLVPEPFDSVTDILARMADAGFSPVEVVWLLSAHSVAAADHVDPTIPGTPFDST
PNLFDSQFFIETQLRGTTFPGTGGNPGEVKSPLPGEMRLQSDHLFARDPRTACEWQSMVNDQQKI
QDRFRDTLFKMSMLGQNQDDMIDCSDVIPVPPPLTTKPHLPAGKSKTDVEQACATAPFPTLPADP
GPPTSVPPVPPA

8H:
AVPPSGKRATCSNGKTVNNDACCVWFDVLDDIQTNLFHGGQCGEDAHEALRLTFHDAIAFSPAL
WAQGQFGGGGADGSIIAHSDIELTYPANNGIDEIVEASRHIAQKHNVSFGDFIQFAGAVGVANCN
GGPQLPFFAGRPNPSQPAPPNLVPLPSDSADQILARFADAGFSAVEVVWLLVSHTVGSQHTVDPSI
PGAPFDSTPSDFDAQFFVETMLNGTLVPGNGLQQGEVNSPYPGEFRLQSDFLLARDPRTACEWQ
KMIADQDNMQSKFAAVMLKMSLLGFDQSSLIDCSDVIPTPPGTVQDPFLPAGLTVDDLQPACSD
SPFPTVPTVPGPATSIPPVPMDS

11H:
VTLPQKRATCSGGQTTSNAACCVLFDLRDDLQKNLFDGGQCGEEVHESLRLTFHDAIGFSPTKG
GGGADGSVLIFSDTELNFPANLGIDEIVEAQKPFLQRHNISAGDLVQFAGALGVSNCPGAPRIPFF
LGRPPATAPSPDGLVPEPFDSVDDILARMADAGFSPVEVVWLLSSHTIAAADHVDPTIPGTPFDST
PSIFDSQFFIETQLRGTLFPGTGGNPGEVESPLPGEIRLQSDHLLARDPRTACEWQSMVDNMPKIQ
NRFAATMLKMSLLGQNVRDLIDCSDVIPTPPPLVGTAHLPAGKTQSDVEQACATTPFPTIPADPG
PVTSVPPVPPS

#### Supplementary Tables

**Table S1.** Selected VPs origins, protein lengths and number of mutations in each design

| Name | Species | Protein length | # mut. H | # mut. M | # mut. L |
|---|---|---|---|---|---|
| VP8 | *Ganoderma* sp. 10597_SS1 | 346 | 38 | 25 | 12 |
| VP2 | *Dichomitus squalens* | 353 | 29 | 18 | 11 |
| VP7 | *Trametes versicolor* | 338 | 49 | 22 | 13 |
| VP4 | *Lentinus tigrinus* ALCF2SS1-6s | 345 | 40 | 21 | 8 |
| VP3 | *Gelatoporia subvermispora B* | 338 | 38 | 21 | 15 |
| VP10 | *Pleurotus* sp. Florida | 344 | 33 | 22 | 11 |
| VPL | *Pleurotus eryngii* (VPL2) | 331 | 39 | 23 | 16 |
| VP5 | *Pleurotus ostreatus* | 339 | 43 | 27 | 18 |
| VP11 | *Pleurotus ostreatus* | 338 | 38 | 23 | 13 |
| VP9 | *Ganoderma* sp. 10597 SS1 | 335 | 30 | 19 | 11 |
| VP1 | *Bjerkandera adusta* | 340 | 27 | 14 | 7 |
| VP6 | *Physisporinus* sp. PF18 | 337 | 25 | 15 | 8 |

### mut refers to the number of mutations in each designed variant: H – high mutational load, M – medium mutational load, L – low mutational load.

**Table S2.** Selected VP sequences homology (in percentage)

| Name | VP8 | VP2 | VP7 | VP4 | VP3 | VP10 | VPL | VP5 | VP11 | VP9 | VP1 | VP6 |
|---|---|---|---|---|---|---|---|---|---|---|---|---|
| VP8 | 100 | 77 | 76 | 75 | 52 | 57 | 57 | 52 | 55 | 53 | 57 | 55 |
| VP2 | | 100 | 80 | 81 | 51 | 58 | 58 | 53 | 58 | 52 | 57 | 55 |
| VP7 | | | 100 | 81 | 53 | 60 | 60 | 55 | 60 | 54 | 60 | 57 |
| VP4 | | | | 100 | 83 | 59 | 59 | 54 | 58 | 55 | 59 | 56 |
| VP3 | | | | | 100 | 64 | 66 | 63 | 66 | 66 | 69 | 69 |
| VP10 | | | | | | 100 | 73 | 66 | 76 | 68 | 68 | 72 |
| VPL | | | | | | | 100 | 69 | 78 | 68 | 70 | 71 |
| VP5 | | | | | | | | 100 | 80 | 61 | 61 | 64 |
| VP11 | | | | | | | | | 100 | 66 | 68 | 72 |
| VP9 | | | | | | | | | | 100 | 71 | 74 |
| VP1 | | | | | | | | | | | 100 | 76 |
| VP6 | | | | | | | | | | | | 100 |

**Table S3.** Most-active VP designs sequences homology (in percentage)

| Name | VPL | 5H | 8H | 11H |
|---|---|---|---|---|
| VPL | 100 | 74 | 66 | 81 |
| 5H | | 100 | 60 | 83 |
| 8H | | | 100 | 64 |
| 11H | | | | 100 |

**Table S4.** Kinetic parameters with various substrates

| Substrate | Kinetic constants | 5H | 8H | 11H | R4 |
|---|---|---|---|---|---|
| ABTS (high efficiency) | $K_M$ (μM)<br>$k_{cat}$ ($sec^{-1}$)<br>$k_{cat}$ / $K_M$ ($sec^{-1}$ $mM^{-1}$) | 1.92 ± 0.37<br>1.47 ± 0.07<br>763 ± 151 | 5.04 ± 0.82<br>7.81 ± 0.28<br>1549 ± 258 | 6.29 ± 0.72<br>1.97 ± 0.07<br>314 ± 37 | 0.39 ± 0.08<br>1.28 ± 0.07<br>3,294 ± 699 |
| ABTS (low efficiency) | $K_M$ (μM)<br>$k_{cat}$ ($sec^{-1}$)<br>$k_{cat}$ / $K_M$ ($sec^{-1}$ $mM^{-1}$) | 765 ± 96<br>33.2 ± 1.8<br>43.4 ± 5.9 | 795 ± 96<br>28.2 ± 0.9<br>35.4 ± 4.4 | 1,084 ± 88<br>11.2 ± 0.3<br>10.34 ± 0.88 | 987 ± 107<br>400 ± 21<br>405 ± 49 |
| DMP (high efficiency) | $K_M$ (μM)<br>$k_{cat}$ ($sec^{-1}$)<br>$k_{cat}$ / $K_M$ ($sec^{-1}$ $mM^{-1}$) | 85.7 ± 16.9<br>1.24 ± 0.08<br>14.4 ± 3.0 | 22.9 ± 1.8<br>2.16 ± 0.08<br>94.6 ± 8.2 | 168.2 ± 9.9<br>1.91 ± 0.04<br>11.4 ± 0.7 | 1.36 ± 0.27<br>0.737 ± 0.050<br>542 ± 113 |
| DMP (low efficiency) | $K_M$ (μM)<br>$k_{cat}$ ($sec^{-1}$)<br>$k_{cat}$ / $K_M$ ($sec^{-1}$ $mM^{-1}$) | 27,672 ± 7,087<br>5.35 ± 0.61<br>0.193 ± 0.054 | 164 ± 12<br>3.62 ± 0.05<br>22.1 ± 1.6 | 11,417 ± 1,991<br>6.34 ± 0.36<br>0.555 ± 0.102 | 16,065 ± 1,684<br>132 ± 7<br>8.19 ± 0.96 |
| $Mn^{2+}$ | $K_M$ (μM)<br>$k_{cat}$ ($sec^{-1}$)<br>$k_{cat}$ / $K_M$ ($sec^{-1}$ $mM^{-1}$) | 3,480 ± 150<br>9.11 ± 0.11<br>2.62 ± 0.12 | 11,300 ± 1,360<br>0.64 ± 0.03<br>0.057 ± 0.007 | 289 ± 34<br>19.68 ± 0.47<br>68.1 ± 8.2 | 523 ± 28<br>78.3 ± 1.1<br>150 ± 8 |
| VA | $K_M$ (μM)<br>$k_{cat}$ ($sec^{-1}$)<br>$k_{cat}$ / $K_M$ ($sec^{-1}$ $mM^{-1}$) | 158 ± 27<br>0.52 ± 0.02<br>3.27 ± 0.57 | 10,070 ± 350<br>5.50 ± 0.08<br>0.547 ± 0.021 | 2,010 ± 140<br>1.38 ± 0.03<br>0.684 ± 0.050 | 7,400 ± 1,040<br>5.37 ± 0.35<br>0.73 ± 0.11 |
| RB5 | $K_M$ (μM)<br>$k_{cat}$ ($sec^{-1}$)<br>$k_{cat}$ / $K_M$ ($sec^{-1}$ $mM^{-1}$) | 2.94 ± 1.04<br>0.18 ± 0.04<br>62.3 ± 25.2 | 17.27 ± 1.74<br>1.12 ± 0.05<br>65.0 ± 7.2 | 1.98 ± 0.24<br>0.30 ± 0.01<br>152 ± 20 | 1.29 ± 0.13<br>0.628 ± 0.020<br>487 ± 51 |
| $H_2O_2$ | $K_M$ (μM)<br>$k_{cat}$ ($sec^{-1}$)<br>$k_{cat}$ / $K_M$ ($sec^{-1}$ $mM^{-1}$) | 197 ± 20<br>20.3 ± 0.8<br>103 ± 11 | 89.7 ± 12.9<br>12.14 ± 0.61<br>135 ± 21 | 38.9 ± 6.9<br>7.01 ± 0.51<br>180 ± 35 | 457 ± 53<br>230 ± 9<br>503 ± 61 |

#### Supplementary Figures

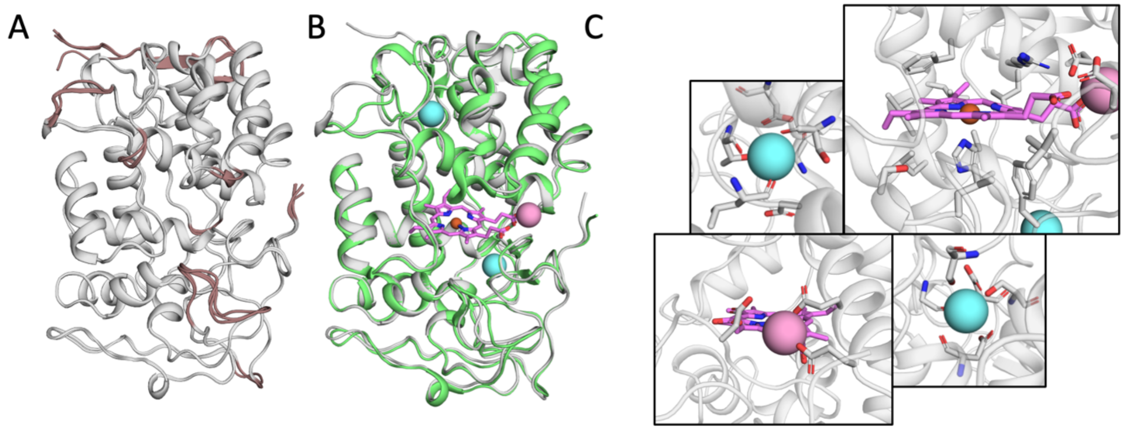


**Figure S1.** Accuracy of trRosetta models. (A) Four most reliable (top-ranked) models of a representative VP (VP5, gray), superimposed onto one another, demonstrate that most of the protein structural elements are converged, and small discrepancies occur only in peripheral loops (brown). (B) Best model of VP5 (gray) overlapped onto wildtype VPL crystal structure (PDB entry 3FJW; light green). VPL calcium and manganese ions are presented in blue and pink spheres, respectively, and the heme group in pink sticks. (C) Close-up look onto VP5 residues that face all ligands and ions.

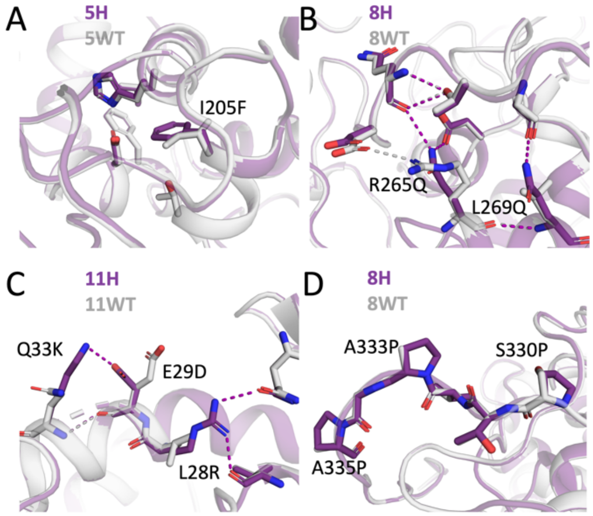


**Figure S2.** Examples of PROSS mutations in VP designs. (A) Ile205Phe in 5H improves core packing. (B) Arg265Gln and Leu269Gln in 8H and (C) Leu28Lys, Glu29Asp and Gln33Lys in 11H generate new hydrogen-bond networks. (D) Mutations at positions 330, 333 and 335 to Pro in 8H improve loop rigidity. In all panels, the PROSS-design model (purple) is superimposed onto the trRosetta-generated wildtype model (gray). Significant mutations, residues in their vicinity and the hydrogen bonds they form are presented in purple and gray sticks for the wildtype and designed models, respectively.

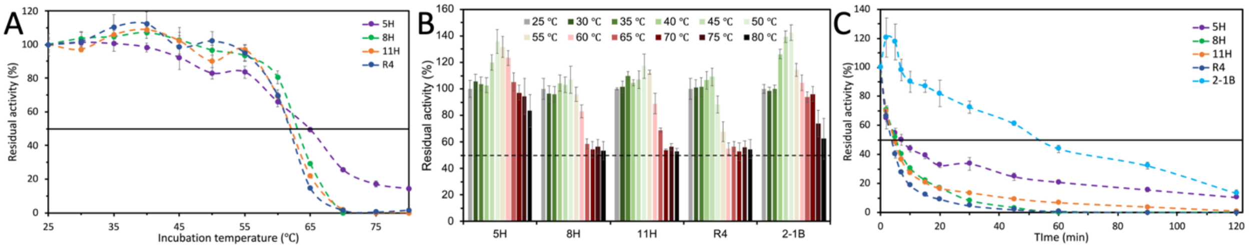


**Figure S3.** Thermal stabilities of selected VP variants (5H, 8H, 11H, VPL-R4 and 2-1B). VPs were incubated for (A) 15 or (B) 10 minutes at temperatures ranging from 30 to 80 °C, and their residual activity compared to the activity at 25 °C was measured. (C) kinetic thermostability ($t_{1/2}$) profiles were determined by incubation of VP supernatants at 65 °C and measuring their residual activity at times 0-120 minutes, compared to the activity at time zero. All the results are the means ± S.D. from three independent experiments.

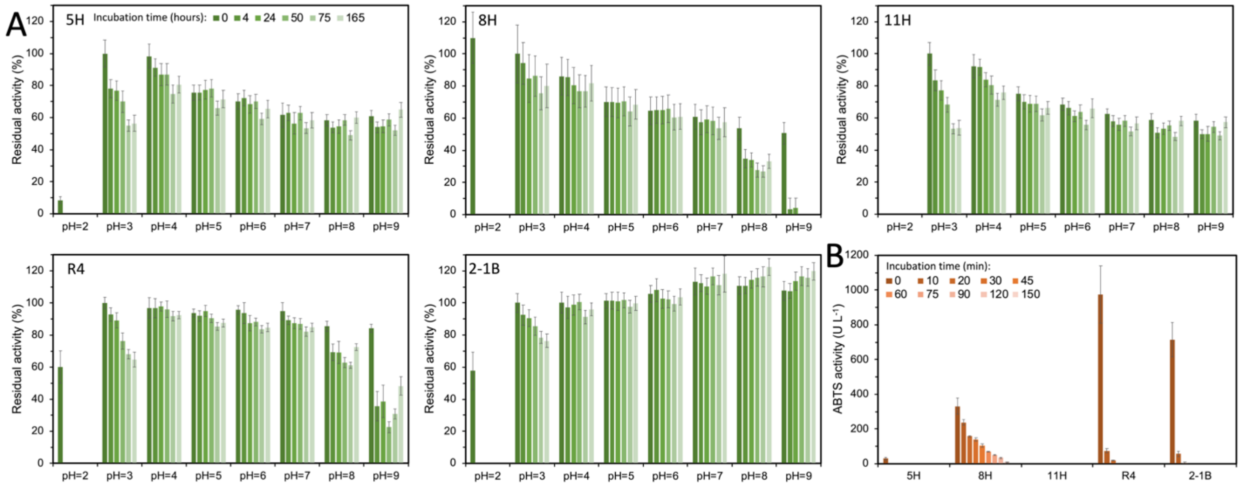


**Figure S4.** pH stabilities of selected VP variants (5H, 8H, 11H, VPL-R4 and 2-1B). (A) VPs were incubated at pH ranging from 2-9 using 100 mM citrate-phosphate-borate buffer, and their residual activity at times 0-165 hours, compared to the activity at pH=3 at time zero, was measured. (B) VPs were incubated at 100 mM citrate-phosphate-borate buffer pH=2, and their activity at times 0-150 minutes was measured. All the results are the means ± S.D. from three independent experiments.

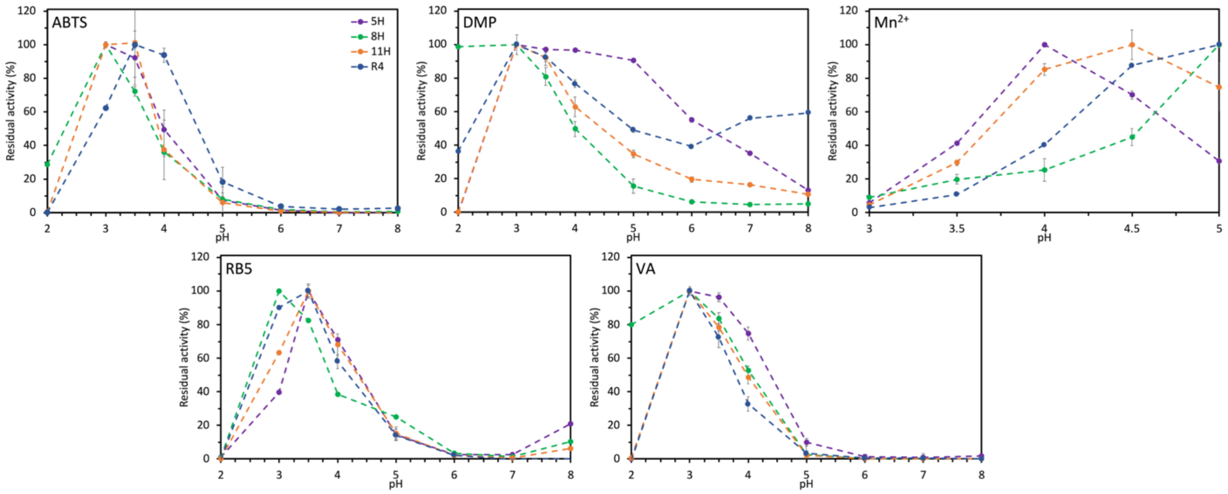


**Figure S5.** pH activity profiles of selected VP variants with versatile substrates. Purified 5H, 8H, 11H and R4 assayed for activity at range of pHs (pHs 2-9 to all substrates but $Mn^{2+}$, in which was tested at pH range of 3-5): ABTS, DMP, $Mn^{2+}$, VA and RB5. VPs activity was normalized to the activity at optimal pH for each protein-substrate pair. All the results are the means ± S.D. from three independent experiments.

#### Supplementary References


(1) Grigoriev, I. V.; Nikitin, R.; Haridas, S.; Kuo, A.; Ohm, R.; Otillar, R.; Riley, R.; Salamov, A.; Zhao, X.; Korzeniewski, F.; Smirnova, T.; Nordberg, H.; Dubchak, I.; Shabalov, I. MycoCosm Portal: Gearing up for 1000 Fungal Genomes. *Nucleic Acids Res.* **2014**, *42* (Database issue), D699–D704.

(2) Savelli, B.; Li, Q.; Webber, M.; Jemmat, A. M.; Robitaille, A.; Zamocky, M.; Mathé, C.; Dunand, C. RedoxiBase: A Database for ROS Homeostasis Regulated Proteins. *Redox Biol* **2019**, *26*, 101247.

(3) Choi, J.; Détry, N.; Kim, K.-T.; Asiegbu, F. O.; Valkonen, J. P. T.; Lee, Y.-H. fPoxDB: Fungal Peroxidase Database for Comparative Genomics. *BMC Microbiol.* **2014**, *14*, 117.

(4) Almagro Armenteros, J. J.; Tsirigos, K. D.; Sønderby, C. K.; Petersen, T. N.; Winther, O.; Brunak, S.; von Heijne, G.; Nielsen, H. SignalP 5.0 Improves Signal Peptide Predictions Using Deep Neural Networks. *Nat. Biotechnol.* **2019**, *37* (4), 420–423.

(5) Edgar, R. C. MUSCLE: Multiple Sequence Alignment with High Accuracy and High Throughput. *Nucleic Acids Res.* **2004**, *32* (5), 1792–1797.

(6) Kumar, S.; Stecher, G.; Li, M.; Knyaz, C.; Tamura, K. MEGA X: Molecular Evolutionary Genetics Analysis across Computing Platforms. *Mol. Biol. Evol.* **2018**, *35* (6), 1547–1549.

(7) Stecher, G.; Tamura, K.; Kumar, S. Molecular Evolutionary Genetics Analysis (MEGA) for macOS. *Mol. Biol. Evol.* **2020**, *37* (4), 1237–1239.

(8) Madeira, F.; Park, Y. M.; Lee, J.; Buso, N.; Gur, T.; Madhusoodanan, N.; Basutkar, P.; Tivey, A. R. N.; Potter, S. C.; Finn, R. D.; Lopez, R. The EMBL-EBI Search and Sequence Analysis Tools APIs in 2019. *Nucleic Acids Res.* **2019**, *47* (W1), W636–W641.

(9) Yang, J.; Anishchenko, I.; Park, H.; Peng, Z.; Ovchinnikov, S.; Baker, D. Improved Protein Structure Prediction Using Predicted Interresidue Orientations. *Proc. Natl. Acad. Sci. U. S. A.* **2020**, *117* (3), 1496–1503.

(10) Hiranuma, N.; Park, H.; Baek, M.; Anishchenko, I.; Dauparas, J.; Baker, D. Improved Protein Structure Refinement Guided by Deep Learning Based Accuracy Estimation. *Nat. Commun.* **2021**, *12* (1), 1340.

(11) Goldenzweig, A.; Goldsmith, M.; Hill, S. E.; Gertman, O.; Laurino, P.; Ashani, Y.; Dym, O.; Unger, T.; Albeck, S.; Prilusky, J.; Lieberman, R. L.; Aharoni, A.; Silman, I.; Sussman, J. L.; Tawfik, D. S.; Fleishman, S. J. Automated Structure- and Sequence-Based Design of Proteins for High Bacterial Expression and Stability. *Mol. Cell* **2016**, *63* (2), 337–346.

(12) Weinstein, J. J.; Goldenzweig, A.; Hoch, S.-Y.; Fleishman, S. J. PROSS 2: A New Server for the Design of Stable and Highly Expressed Protein Variants. *Bioinformatics* **2020**. https://doi.org/10.1093/bioinformatics/btaa1071.

(13) Jumper, J.; Evans, R.; Pritzel, A.; Green, T.; Figurnov, M.; Ronneberger, O.; Tunyasuvunakool, K.; Bates, R.; Žídek, A.; Potapenko, A.; Bridgland, A.; Meyer, C.; Kohl, S. A. A.; Ballard, A. J.; Cowie, A.; Romera-Paredes, B.; Nikolov, S.; Jain, R.; Adler, J.; Back, T.; Petersen, S.; Reiman, D.; Clancy, E.; Zielinski, M.; Steinegger, M.; Pacholska, M.; Berghammer, T.; Bodenstein, S.; Silver, D.; Vinyals, O.; Senior, A. W.; Kavukcuoglu, K.; Kohli, P.; Hassabis, D. Highly Accurate Protein Structure Prediction with AlphaFold. *Nature* **2021**, *596* (7873), 583–589.

(14) Alcalde, M. Mutagenesis Protocols in Saccharomyces Cerevisiae by In Vivo Overlap Extension. In *In Vitro Mutagenesis Protocols: Third Edition*; Braman, J., Ed.; Humana Press: Totowa, NJ, 2010; pp 3–14.

(15) Viña-Gonzalez, J.; Gonzalez-Perez, D.; Ferreira, P.; Martinez, A. T.; Alcalde, M. Focused Directed Evolution of Aryl-Alcohol Oxidase in Saccharomyces Cerevisiae by Using Chimeric Signal Peptides. *Appl. Environ. Microbiol.* **2015**, *81* (18), 6451–6462.

(16) Garcia-Ruiz, E.; Gonzalez-Perez, D.; Ruiz-Dueñas, F. J.; Martínez, A. T.; Alcalde, M. Directed Evolution of a Temperature-, Peroxide- and Alkaline pH-Tolerant Versatile Peroxidase. *Biochem. J* **2012**, *441* (1), 487–498.
